# Use of Genome Scale Metabolic Reconstructions of Avian Pathogenic *Escherichia coli* (APEC) phylogroups for the identification of lineage-specific metabolic pathways

**DOI:** 10.1101/2024.12.20.629819

**Authors:** Huijun Long, Jai W. Mehat, HuiHai Wu, Arnoud H. M. van Vliet, Roberto M. La Ragione

## Abstract

Avian Pathogenic *Escherichia coli* (APEC) are a genetically diverse pathotype primarily associated with extra-intestinal infections in birds. APEC lineages are predicted to have unique metabolic capabilities contributing to virulence and survival in the host environment. Here we present a genome-scale metabolic model for the APEC pathotype based on 114 APEC genome sequences, and lineage-specific models for the phylogroups B2, C and G based on a representative isolate for each phylogroup. A total of 1,848 metabolic reactions were predicted in the 114 APEC isolates before gap filling and manual correction. Of these, 89% represented core reactions, whilst the 11% accessory reactions were mostly associated with carbon and nitrogen metabolism. Predictions of auxotrophy were confirmed by inactivation of the conditionally essential *lysA* and the non-essential *potE* genes. The APEC metabolic model outperformed the *E. coli* K-12 *i*JO1366 model in the Biolog Phenotypic Array platform. Sub-models specific for phylogroups B2, C and G predicted differences in the metabolism of 3-hydroxyphenylacetate (3-HPAA), a phenolic acid derived from the flavonoid quercetin, which is commonly added to poultry feed. Two 3-HPAA associated reactions/genes distinguished APEC phylogroup C from APEC phylogroups B2 and G, and 3-HPAA supported the growth of APEC phylogroup C in minimal media, but not phylogroups B2 and G. In conclusion, we have constructed genome-scale metabolic models for the three major APEC phylogroups B2, C and G, and have identified a metabolic pathway distinguishing phylogroup C APEC. This demonstrates the importance of lineage- and pathotype-specific metabolic models when investigating genetically diverse microbial pathogens.

**IMPACT STATEMENT:** Avian Pathogenic *Escherichia coli* (APEC) are the cause of colibacillosis in poultry, which results in a significant economic burden to the poultry industry, and strongly affects the health and welfare of flocks. APEC isolates show a high level of genetic diversity, which complicates diagnostics, epidemiology and the design of prevention and treatment strategies. In this study we have used genome sequences derived from 114 APEC isolates to investigate their metabolic capabilities, and define the metabolic diversity of APEC within a generalised APEC metabolic model, and lineage-specific metabolic models. These models have been interrogated to find unique pathways that can be targeted for the development of anti-APEC treatments, and one such metabolic pathway was identified as a proof of principle. This approach shows great promise for the design of future strategies to prevent and deal with APEC infections, and can be adapted to other genetically diverse microbial pathogens.

## INTRODUCTION

Avian pathogenic *Escherichia coli* (APEC) belong to the extra-intestinal pathogenic (ExPEC) group of *E. coli* [1, 2]. APEC cause extra-intestinal infections in birds, commonly termed colibacillosis, a collective name describing a range of systemic and localised syndromes such as pericarditis, perihepatitis, peritonitis, and airsacculitis. Colibacillosis causes significant economic losses and is a major welfare concern in the poultry industry [3]. The APEC pathotype is defined by the diseases caused in birds, with *E. coli* isolated from extra-intestinal locations in birds with colibacillosis-associated diseases being candidate APEC, pending further characterisation. APEC are found in all major *E. coli* lineages and phylogroups and encompass a range of serogroups and MLST sequence types [4–7], and it this genetic variation and imprecise definition which complicates the development and efficacy of control strategies, such as antimicrobial administration and vaccine development [6]. Therefore, an improved understanding of the biology and variation of this pathogen will be beneficial to the development of novel and effective control strategies.

It has been proposed that different pathogenic *E. coli* lineages, in addition to commensal lineages, harbour variable metabolic profiles [8–10]. However, there remains a paucity of knowledge regarding the unique metabolic traits of APEC isolates, as most studies have focused on identification of markers allowing distinction of APEC from avian faecal *E. coli* [7]. Genome scale metabolic reconstructions (GEMs) are a powerful tool that have been applied in a number of fields [11]. Essentially, GEMs use genome sequences and database comparisons to predict metabolic functions and responses in a range of environmental conditions. To date, most *E. coli* GEMs have been constructed from a single commensal isolate, such as K-12 MG1655 [12–15], probiotic isolate Nissle 1917 [16] and *E. coli* W (ATCC 9637) [17]. However, as increasing numbers of *E. coli* genome sequences are now available, it has become apparent that GEMs of single commensal isolates only partially represent the metabolic capability of this species and its different pathotypes [9]. Similarly, it was reported that GEMs based on multiple isolates were able to differentiate the metabolic profile of pathogenic from commensal *E. coli* isolates [9]. Hence it is clear that single laboratory strain-based models are not adequate to represent the metabolic capability and variation within *E. coli* pathotypes.

In this study, we have constructed a comprehensive metabolic genome scale model based on a panel of 114 APEC isolates. This general APEC model was then used as a template to develop specific models that recapitulate the metabolism of the major phylogroups of the APEC pathotype. Models were queried for auxotrophy and gene essentiality, growth predictions in diverse media and used to identify a metabolic pathway distinguishing phylogroup C APEC from phylogroup B2 and G APEC, demonstrating the potential for such GEMs to improve our understanding of APEC biology and identification of potential targets for future prevention and intervention strategies.

## METHODS

### Bacterial strains and growth conditions

114 APEC isolates were obtained from the SAP culture collection at the School of Veterinary Medicine, University of Surrey and used for APEC model construction (Supplementary Material 2, Table S1). These APEC isolates were obtained from commercial laying hens or broilers with either colibacillosis, perihepatitis, yolk sac infection, pericarditis or peritonitis in the UK and Germany. Isolates were collected at *post-mortem* examination per the protocols used at the diagnostic services involved, and were submitted to the School of Veterinary Medicine at the University of Surrey between 2014 and 2017 for further diagnostic or research activity. The clinical information available is provided in Supplementary Material 2, Table S1. All isolates were stored in BHI broth supplemented with 50% glycerol at -80 °C until required for analysis. Prior to analysis, the APEC isolates were streaked onto MacConkey agar 3 plates, Nutrient agar or Luria-Bertani (LB) and incubated aerobically at 37°C for 16-24 hours, unless otherwise stated. M9 minimal medium (1× M9 salt, 2 mM MgSO4, 0.1 mM CaCl2, and 0.4% glucose if required) was used for specific experiments as indicated below. Chloramphenicol and ampicillin were used at final concentrations of 100 µg/ml and 30 µg/ml, respectively. All growth media and antibiotic supplements were purchased from Oxoid (ThermoFisher Scientific) unless otherwise stated.

### Biolog Phenotypic Microarray and M9 minimal medium test supplemented with 3-hydroxyphenylacetate (3-HPAA)

Three APEC isolates were selected from the representative phylogroups B2, C and G (serogroups O2, O78, and O24, respectively) and used for Biolog analysis (N=3). Additional isolates from these phylogroups were selected for M9 minimal medium cultivation analysis (N=13). The isolates used for these analyses are listed in Supplementary Material 2, Table S1. Three biological and three technical replicates were performed for each isolate. The APEC isolates were pre-cultured on R2A agar plates (Biolog) for Biolog assays as per the manufacturer’s instructions [18], and subsequently tested using 190 carbon sources (PM1 and PM2 MicroPlate^TM^), as well as 59 Phosphorus and 35 Sulphur Sources (PM4A MicroPlate^TM^). The cell suspensions and plate inoculation were undertaken according to the instructions from the manufacturer. The Biolog PM1, PM2 and PM4A microplates were incubated and analysed in Biolog’s OmniLog® instrument at 37 °C for 24h, aerobically.

Readings were taken every 15 min. M9 minimal media was used to test the metabolic ability of isolates in a defined synthetic medium. Three concentrations (1 mM, 0.75 mM, and 0.5 mM) of 3-hydroxyphenylacetate (3-HPAA) were tested in this study. L-lysine or putrescine (1 mM) was also tested in this study for the cultivation analysis of gene knockout mutants. The polystyrene 96 well plates (ThermoFisher Scientific) were cultured and read in Spark® microplate reader (TECAN) at 37 °C for 24h, aerobically. Readings were taken every 15 min.

### DNA extraction and Next generation sequencing (NGS)

The high molecular weight genomic DNA of APEC isolates was extracted and purified using a DNA Wizard® genomic purification kit (Promega), according to the instructions from the manufacturer. The extracted DNA samples were sent to the Animal and Plant Health Agency (Weybridge, UK) for sequencing on the Illumina MiSeq platform, providing 150 nt paired end reads. The obtained raw DNA sequences were *de novo* assembled in contigs using Shovill version 1.1.0 (https://github.com/tseemann/shovill) using version 3.14 of the Spades assembler [19] with minimum coverage of 2.00× and minimum length of 200 nt. Quality assessment of genome assemblies were taken by QUAST version 5.0 [20], and required the following minimum metrics to be satisfied: genome size between 4.5-6.0 Mbp, number of contigs < 500 and N_50_ values of > 50 kb, respectively (Supplementary Material 2, Table S1). Prokka version 1.14 [21] was used for initial annotation of genome sequences, which produced relevant output files compatible for downstream analyses. The phylogenetic grouping was analysed on the *in-silico* based ClermonTyping program [22]. *In silico* serotyping was performed using Abricate version 1.0.1 (https://github.com/tseemann/abricate) with the EcOH [23] database. MLST sequence typing was performed using the mlst v2.23 program (https://github.com/tseemann/mlst) with the Warwick scheme [24]. A phylogenetic tree based on whole genome single-nucleotide polymorphisms (SNPs) was generated using ParSNP version 1.2 [25], resulting in a phylogenetic tree highlighting genomic similarity and sharing of core genome of all isolates. All 114 APEC genomes were screened for 11 proposed virulence markers [26] using Abricate. All relevant information for the 114 genomes (assembly metrics, presence/absence of virulence genes, O:H serotype information and MLST sequence types) are included in Supplementary Material 2, Table S1.

### Construction of APEC metabolic models

The FASTA files of 114 APEC samples generated from Prokka were submitted to the RAST server (https://rast.nmpdr.org/) for genome annotation [27]. The RAST annotation was automatically submitted to ModelSEED (https://modelseed.org/genomes/) for further model draft construction [28]. Model gap-filling is an essential step to investigate the missing reactions, which apply a mixed-integer linear program to define the minimal number of reactions required for the customer-defined collection of reactions. Several algorithms have been reported and applied in model gap-filling [29, 30]. COBRA Toolbox v3.0 [31] was used in this study and the database was based on ModelSeed. Further manual curation was performed when model was unable to be corrected by gap-filling steps, for instance: unnecessary reactions must be on for model running, and to avoid stoichiometrically balanced cycles which are a subset of contiguous reactions that forms a loop or internal network reactions, that carry fluxes without any exchange reactions involved. The model was adjusted based on the literature (basic nutrient requirement for cell growth and common pathways) and reference model *i*JO1366 [14]. The correction procedure included adding/deletion of reactions, adjusting of reversibility/directionality of reactions, and change of biomass consumption reactions. Metabolic models have been included as Supplementary Material 3 (Table S5), Supplementary Material 4 (Table S6) and Supplementary Material 5 (Table S7).

### GEM analysis

The growth prediction analysis was performed on the SurreyFBA 2.0 beta with JyMet2 [32]. The model was imported as a SurreyFBA formatted file, meanwhile a specific format (pfile) was also introduced, in which the media setting can be adjusted via changing the upper/lower bound of each exchange reaction. The computational prediction of growth rate was based on typical flux units (millimole per gram dry weight per hour, mmol gDW^−1^ h^−1^). The default nutrient sources were glucose (carbon, C), ammonia (NH_3_), phosphate (PO_4_^2-^) and sulphate (SO_4_^2-^). When examining the usage of a particular nutrient source, the exchange reaction of its corresponding default source was adjusted to zero (reaction shut down). In addition, growth simulation studies were performed using the *E. coli* K-12 model *i*JO1366 [14] for comparative analysis. Due to the differences in default nutrient requirements in modelling, two growth simulations (*i*JO1366a and *i*JO1366b) were performed within two different input nutrient settings. The input nutrients of *i*JO1366a were adjusted as in the APEC model used here, where the exchange reactions of selenate (SeO_4_^2-^), selenite (SeO_3_^2-^) and tungstate (WO_4_^2-^) were disabled as they are absent from the APEC model; whilst these three exchange reactions were enabled in *iJO1366*b to perform the optimal conditions for *i*JO1366. In both *i*JO1366a and *i*JO1366b, the exchange reactions of molybdate (MoO_4_^2-^) and nickel (Ni^2+^), which are also absent from the APEC model, were kept enabled, as they are essential for *i*JO1366. The phenotypic and computational measurements were compared using a Pearson correlation coefficient test to measure the strength and direction of these two variables.

### Inactivation of genes in *E. coli*

Genes were inactivated using the Red Lambda Recombination assay [33]. Two gene targets (*lysA* and *potE*) were selected based on gene essentially predicted by model. The target genes were disrupted by deletion of approximately 450∼550 bp sequence in the middle part of the gene sequence, and replaced with a chloramphenicol resistance cassette (CAT) as selective marker. These cloning fragments were constructed using gBlocks gene fragments in tubes service (gBlocks™, Integrated DNA Technologies). APEC isolates were selected for gene inactivation experiments from phylogroup B2, G and C, respectively, with 90% coverage and identity for the *lysA* and *potE* genes, and no resistance to ampicillin and chloramphenicol in Mueller Hinton broth, as these were used as selective markers for the inactivation procedure and associated plasmids. Electrocompetent cells were generated by three times washing with ice-cold 10% glycerol and electroporated with the assistant plasmid pSIJ8 [33, 34], containing an ampicillin resistance gene. The transformed cells with pSIJ8 were selected on LB agar supplemented with ampicillin, followed by PCR using primers described in Supplementary Material 2, Table S8. The expression of the pSIJ8-encoded λ-Red recombination system was induced with 0.5M arabinose. The cloning fragment was then introduced through electroporation, with cells recovered at 30°C (as plasmid pSIJ8 is temperature sensitive) on a shaking platform for 1 -2h. Mutants were selected on chloramphenicol-containing LB plates and verified by PCR. We also designed a reverse transcription quantitative-PCR (RT-qPCR) to confirm the absence of *lysA* and *potE* mRNA transcripts in mutant candidates. The primers and probe of the real time quantitative PCR were designed to amplify the sequences deleted from the wildtype isolate during the construction of the cloning fragment (Supplementary Material 2, Table S8).

## RESULTS

### Genomic characterisation of APEC isolates for metabolic model generation

A total of 114 *E. coli* isolates were included in this study, isolated from diseased birds from the UK and Germany. The pathologies observed are provided in Supplementary Material 1, Table S1, and ranged from colibacillosis, perihepatitis, peritonitis, pericarditis to yolk sac infection. To test whether these 114 APEC isolates were representative of the APEC pathotype, their genome sequences were determined using Illumina sequencing and subsequent genome assembly. The total size of the genome assemblies was between 4.72-5.56 Mbp (average 5.11 Mbp), with the number of contigs ranging from 104-327 (average 197), and the N_50_ ranging from 63-342 kbp (average 159 kbp). Subsequently their phylogroups, dominant serogroups and sequence types were determined using the genome sequences (Figure 1). The phylogenetic grouping of isolates based on core genome SNPs matched the distinctions in phylogroups and sequence types. *In silico* phylogroup typing showed that the majority of APEC isolates belonged to B2 (27.2%), followed by C (18.4%), G (16.7%), A (15.8%) and B1 (13.2%). The phylogroups D, E and F were underrepresented in this collection. The most prevalent serogroups were O78 (17%) and O2 (13%) and the most common MLST sequence types (STs) were ST-117 (15.8%), ST-23 (12.3%) and ST-95 (10.5%). Screening of the genome sequences for the presence of 11 virulence markers showed that 83/114 (73%) of the included APEC isolates had 5 or more of these virulence markers. All these characteristics broadly matched earlier studies [4–7, 35], and hence we consider this collection of isolates broadly representative for the APEC pathotype.

**Figure 1.**
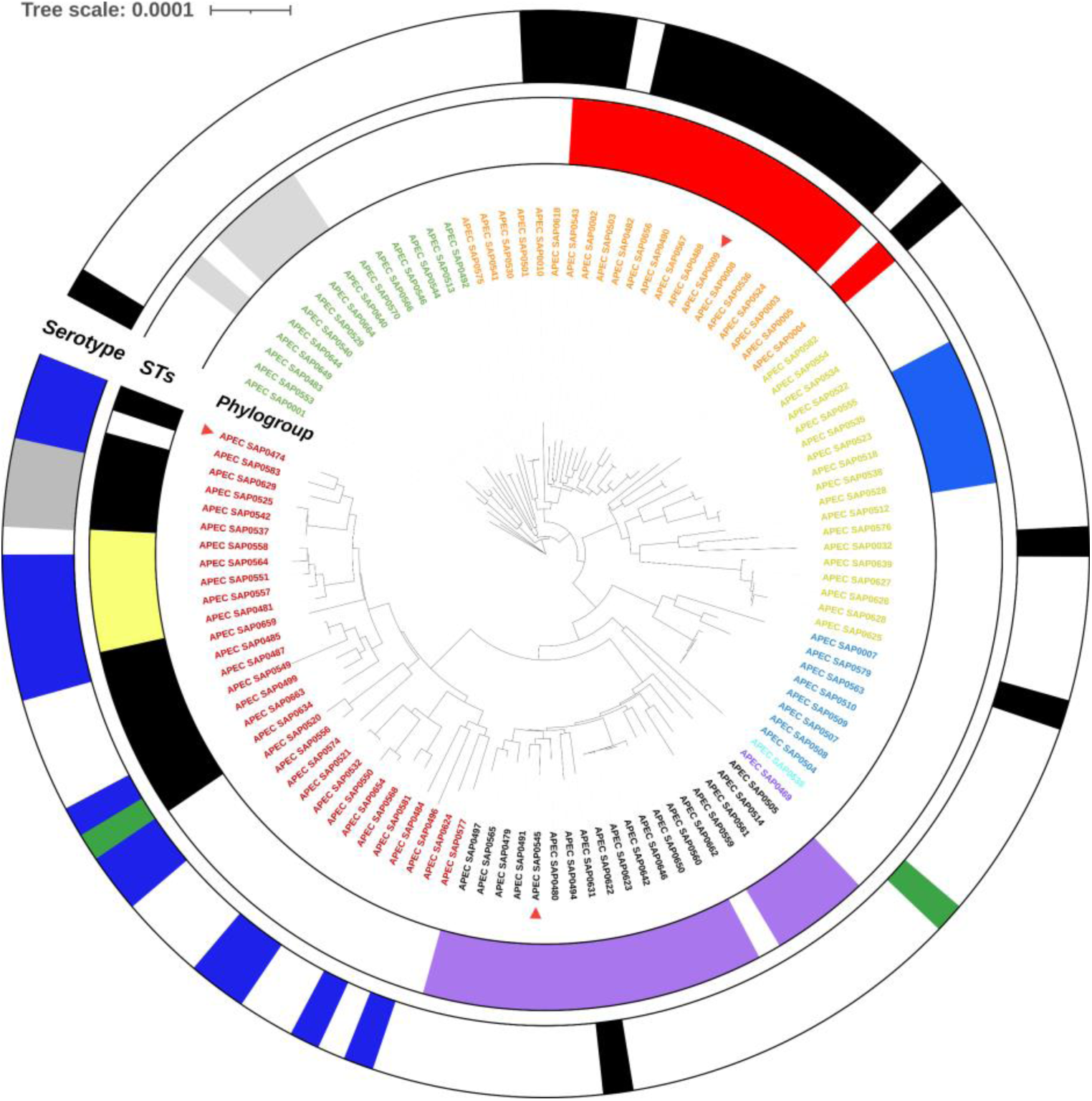
APEC population structure for the isolates selected for GEM construction in this study assigned with phylogroup (n=114). Colour code of phylogroups: A = Yellow, B1 = Green, B2 = Red, C = Orange, D = Purple, E = Blue, F = Light blue, G = Black. The distribution of dominant serogroups is shown in the outer ring. The colour code of serogroup: O78 = Black, O2 = Blue, O18ac = Grey, O1 = Green. The distribution of dominant Sequence types (STs) is shown in the middle ring. The colour code for sequence types: ST-117 = Purple, ST-23 = Red, ST-95 = Black, ST-93 = Blue, ST-101 = Grey, ST-140 = Yellow. The three representative APEC isolates from phylogroup B2, C and G used for Biolog phenotype assays are highlighted with a red triangle symbol.

### Generation of the metabolic profile of the APEC pathotype

The metabolic profile of the APEC isolates was predicted using ModelSEED. A total of 1,848 different metabolic reactions were identified and annotated in the 114 APEC isolates included, and were defined as pan reactions. A total of 1,639 reactions (88.7%) were shared among all isolates, defined as core reactions, whereas 209 reactions (11.3%) were defined as accessory reactions. The reactions were categorised according to their predicted metabolic functions using ModelSEED, as shown in Figure 2. The most highly conserved metabolic reactions were responsible for lipid metabolism (96.4%), nucleotide metabolism (95.1%), coenzyme metabolism (94.2%) and energy production (93.9%). Reactions for alternative carbohydrate utilisation and amino acid metabolism made up 42.9% (792/1,848) of pan reactions and 52.6% (110/209) of accessory reactions. The metabolic reactions were subsequently clustered with the phylogroups identified (Figure 3, Supplementary Material 2, Table S9). In the accessory reactions, there was clustering of metabolic profiles of APEC isolates from the same phylogroups, especially in metabolism of alternative carbohydrates and amino acids including their transport/exchange capabilities. This suggests that certain lineages are likely adapted to niches or environments, where they may harbour an advantage in the utilisation of available nutrient. Notably, phylogroup B2 APEC isolates displayed distinct metabolic capabilities compared to the other phylogroups. Similar observations were also reported previously [36, 37]. Metabolic capabilities absent in phylogroup B2 were mainly associated with amino acids and carbohydrates, and related transport/exchange of metabolites. In contrast, metabolic reactions involved in membrane biogenesis, such as N-acetylneuraminic acid (Neu5Ac) metabolism, were predominantly clustered in phylogroup B2 and G. These reactions have been described to contribute to the pathogenicity of Gram-negative bacteria by protecting against the host immune system [38].

**Figure 2.**
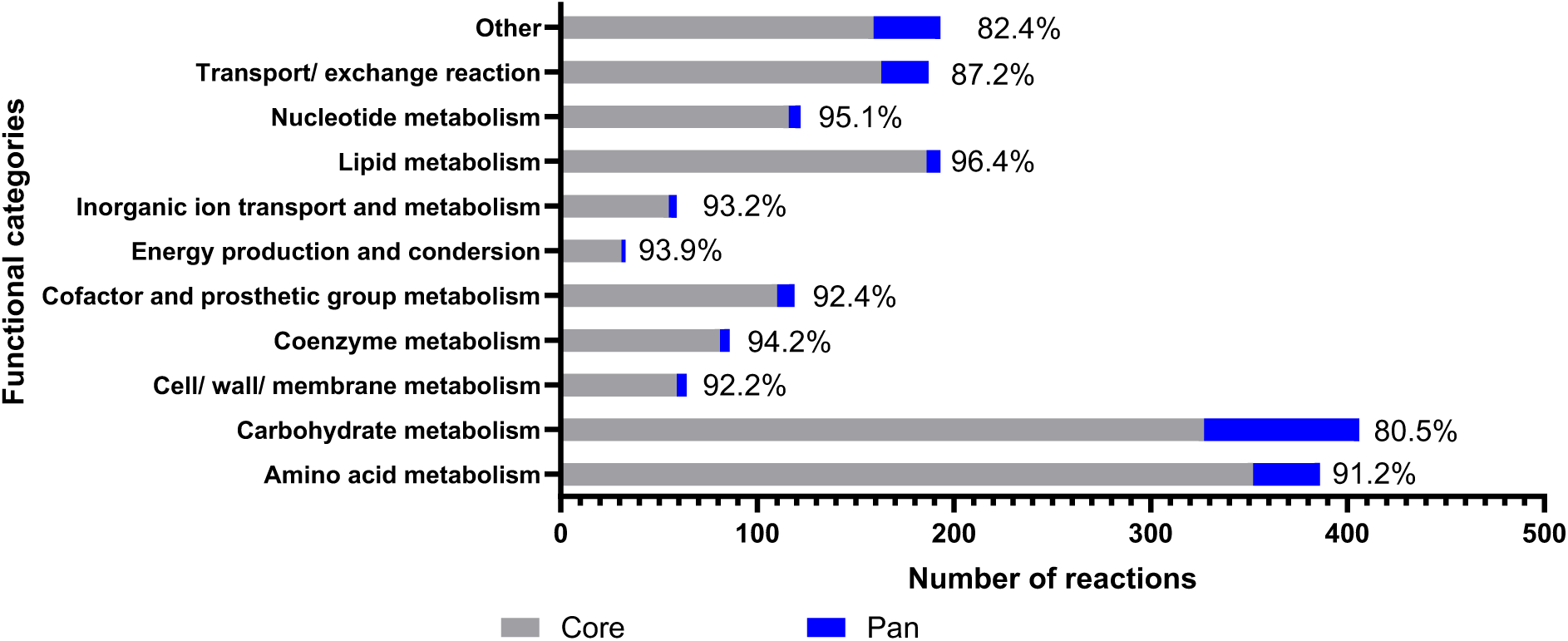
The distribution of annotated metabolic reactions screened by ModelSEED in 114 APEC isolates. This figure illustrates the core (Grey) and pan (Blue) metabolic content of the 114 APEC isolates used in this study. The core and pan content were defined by the extent of content shared among all isolates. The percentage shown represents the core reactions of the total.

**Figure 3.**
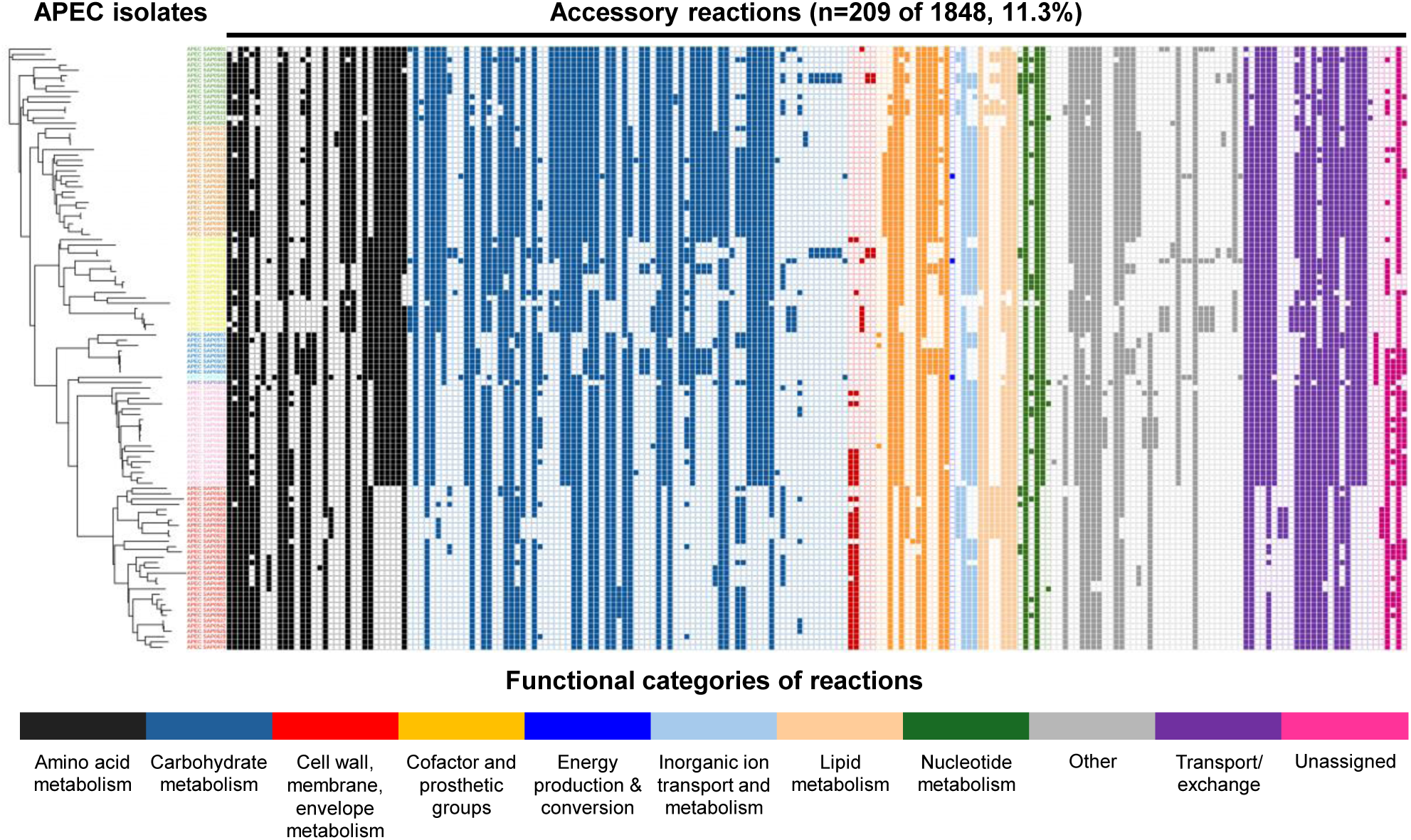
Distribution of accessory metabolic reactions in APEC (n=114) among different phylogenetic groups, clustered using a tree based on single nucleotide polymorphisms. Colour code of phylogroups (from top to bottom): Yellow = A; Green = B1; Orange = C; Dark blue = E; Purple = D; Light blue = F; Red = B2; Pink = G. Presence of metabolic reactions is represented using a filled square, absence of the reaction by an open square. Only the 209 accessory reactions of the total of 1,848 reactions (pan) are shown here; a full overview of reactions and distribution in the 114 APEC is provided in Supplementary Material 2, Table S9. The functional categories of accessory reactions is also shown here. Colour code of functional categories of reactions (from left to right): Black = Amino acid metabolism; Blue = Carbohydrate metabolism; Red = Cell wall/ membrane/ envelope metabolism; Orange = Cofactor and prosthetic group metabolism; Dark blue = Energy production and conversion; Light blue = Inorganic ion transport and metabolism; Beige = Lipid metabolism; Green = Nucleotide metabolism; Gray = Other; Purple = Transport/exchange reaction; Dark pink = unassigned (* Unassigned reactions were not involved in reconstruction of APEC GEM or phylogroup-specific Sub-GEMs, as determined as dead-end reactions by gap-filling).

### Characterisation of APEC GEM and phylogroup-specific sub-models

The 1,848 metabolic reactions identified in the APEC isolates were used to generate a draft genome-scale metabolic model using ModelSEED. Sixteen reactions were excluded as dead-end reactions. Eventually, the pan-genome metabolic model constructed in this study, referred as APEC GEM, was composed of 2,923 reactions (inclusive of gap-filling reactions and a total biomass reaction), 2,527 metabolites, and 2,242 genes (Figure 4). Notably, there were many reactions added during gap-filling algorithms and manual curation, such as spontaneous reaction for extracellular metabolites exchange, isomerization, putative reactions, which significantly change the distribution of functional categories compared to Figure 2. Based on KEGG annotations and the *E. coli* K-12 model [14], a large number of exchange reactions of extracellular metabolites were accumulated in the transport/exchange functional category (Figure 4). This contrasted with the central metabolism reactions, such as those involved in energy production and nucleotide metabolism, which are generally conserved across *E. coli* lineages.

**Figure 4.**
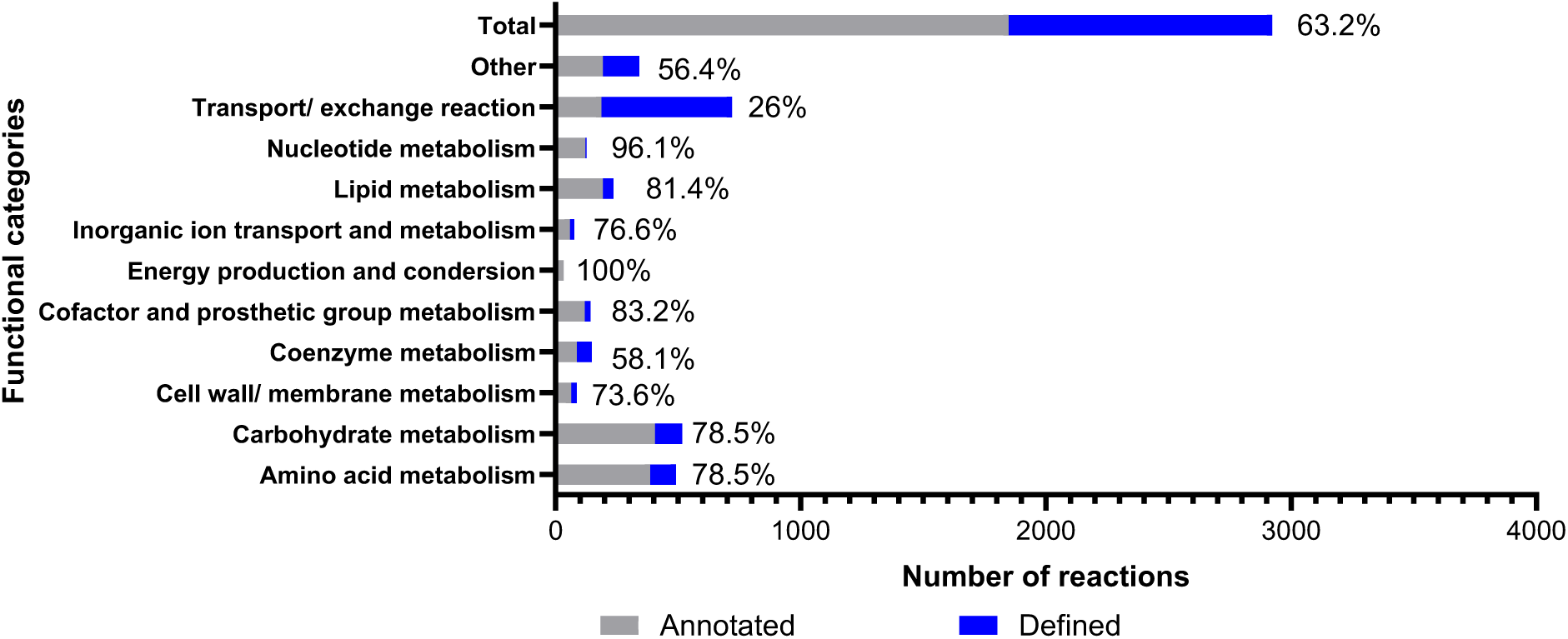
The distribution of the final defined reactions (Grey) in the APEC model after gap-filling and manual curation compared to the annotated metabolic reactions (Blue) screened by ModelSEED in 114 APEC isolates. The percentage shown represents the fraction of reactions retained in the model.

The APEC GEM was built as a pan-genome metabolic model, to represent the full metabolic capabilities in the complete APEC collection used in this study. Considering the distinct metabolic variances between different APEC lineages (Figure 3, Supplementary Material 2, Table S9), phylogroup B2, C and G-specific models were generated from the APEC GEM to represent the metabolic capability and deficiency of each phylogroup. A brief summary of the reactions associated with different metabolic categories removed for sub-model construction for each phylogroup is shown in Supplementary Material 1, Figure S1. In total, 133 reactions were deleted during the construction of sub-models. Notably, the same set of metabolic pathways were removed from more than one phylogroup. Most of deleted reactions were associated with carbohydrate metabolism, accounting for 42%, followed by amino acid metabolism (14.3%) and transport/exchange reactions (9%). A relatively high ratio of carbohydrate (13/19) and amino acid (45/57) metabolism reactions and transport/exchange reactions (9/12) were removed in phylogroup B2 in this study, consistent with prior observations. Besides, 39/57 of carbohydrate metabolism reactions were removed for phylogroup G.

### Gene essentiality analysis of APEC GEM

Gene essentiality analysis is often adopted to evaluate the validity of GEM by comparing with experimentally obtained essential gene datasets [9, 16]. The APEC GEM was used to predict growth on glucose and glycerol as carbon sources. For growth on glucose or glycerol, the lower boundaries of the glucose and glycerol exchange reactions were set to -1,000, respectively. All 2,242 genes in APEC GEM were inactivated *in silico* individually as time and growth were simulated by flux balance analysis (FBA) (Supplementary Material 2, Table S2). When inactivation of a gene resulted in a growth rate above zero, it was considered as a non-essential gene. The gene essentiality predictions on glucose and glycerol in minimal media showed a consistent outcome using the APEC model in this study (Supplementary Material 2, Table S2). In general, *in silico* inactivation of 208 out of 2,242 genes (9.3%) led to an absence of flux through the network and as such would lead to growth inhibition, and thus were determined to be essential genes in the APEC model. The distribution of these essential genes is shown in Supplementary Material 1, Figure S2. Of these essential genes, 57 (27%) were associated with more than one reaction in this model. 13% were responsible for amino acid metabolism and 3% for carbohydrate metabolism. To validate these predictions, we decided against the use of the Keio collection grown on glucose/glycerol, as there are significant differences between *E. coli* K-12 and APEC. Hence two single gene knockout mutants were constructed in APEC for this purpose, in the essential *lysA* gene and non-essential *potE* gene. Both genes were present in 100% of the 114 APEC isolates analysed in this study.

The *lysA* gene is predicted to encode a diaminopimelate decarboxylase that is responsible for the biosynthesis of lysine. In the model, the *lysA* gene (model ID: peg18487) was determined to be an essential gene and encoded for chemical equation (model ID: R_rxn00313) as below ( [c] represent for the metabolites in cytoplasm):

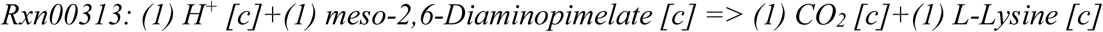

The *potE* gene encodes a membrane protein mediating both the uptake (via proton symport mechanism) and excretion of putrescine (as a putrescine and ornithine antiporter). There is redundancy in putrescine transport with the PotABCD, PotFGHI, PlaP/YeeF, PuuP and SapBCDF systems [39–42]. In the model, the target gene *potE* (model ID: peg6625) was determined as non-essential and responsible for two putrescine transport reactions (model ID: R_rxn05867 and R_rxn10182). For the modelling purpose, the chemical equations of these two reactions are represented as below ( [e] represents for the extracellular metabolites; [c] represent for the metabolites in cytoplasm):

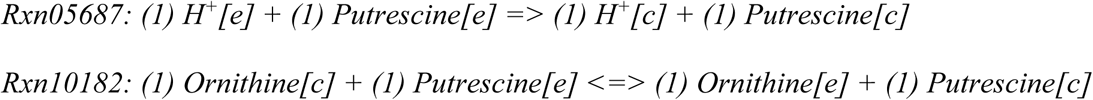

The *lysA* mutants generated in isolates SAP0010 and SAP0631 and *potE* mutant generated in isolates SAP0631 were analysed using glucoside as default carbon source. Other default nutrient sources were adjusted by general standard principle described in next section. The *in-silico* growth prediction of knockouts grown in with and without supplementation of L-lysine or putrescine were compared to the phenotypic screening using M9 minimal medium as illustrated in Figure 5. The Δ*lysA* mutants (Δ*lysA* 10-9 and Δ*lysA* 631-11) were observed to be auxotrophic for L-lysine (Figure 5), which indicates the fact that the *lysA* gene encodes the enzyme in the final and only lysine biosynthesis pathway. This was also revealed in APEC GEM knockout analysis that no biomass was produced when the rxc00313 was switched off. On the contrary, the supplementation of putrescine in M9 medium did not lead any significant change in growth between the Δ*potE* 631-12 mutant and its parental strain (Figure 5). The APEC GEMs correctly predicted the growth status of these knockout mutants, as well as their wildtype, in minimal media with and without supplementation.

**Figure 5.**
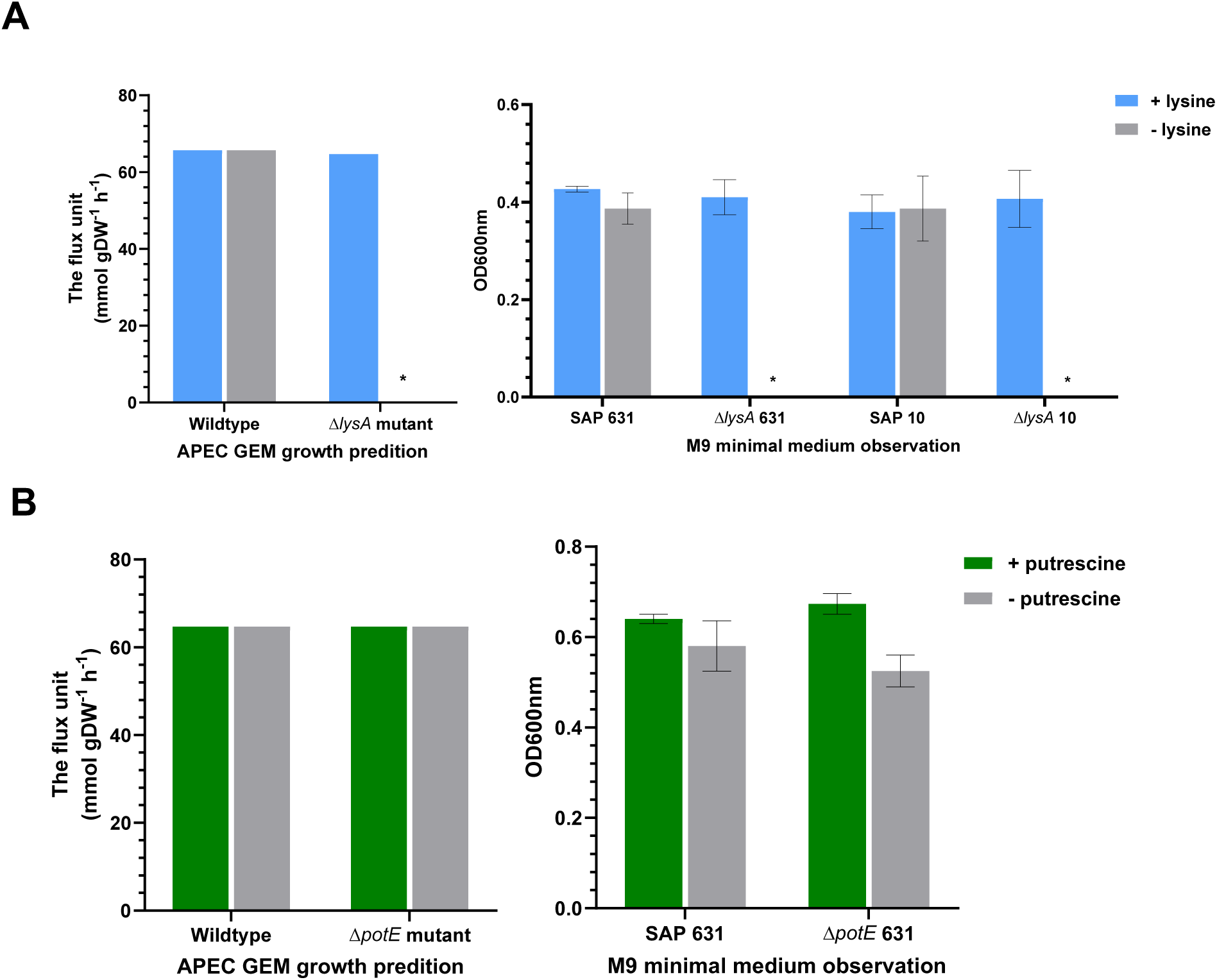
The computational growth prediction (Left) and phenotypic observation (Right) of wildtype and single gene knockout isolates under constrained nutrient input. Figure 5A shows the effect of inactivation of *lysA* with/without medium supplementation with lysine; an asterisk indicates lack of growth demonstrating the essential role of the *lysA* gene. Figure 5B shows the effect of inactivation of *potE* with/without medium supplementation with putrescine, which had no effect as the gene is not essential. The computational prediction is generated by APEC GEM and data is measured by flux unit (mmol gDW^−1^ h^−1^); The phenotypic observation is measured using M9 minimal medium and the optical density is recorded after 24 hour incubation (37 °C) at a wavelength of 600 nm in 96-well plate. The error bar shows standard deviation (SD) of the mean of 3 biological replicates. Asterisks indicate an absence of growth.

### Comparison of the APEC GEM with the *E. coli* K-12 *i*JO1366 GEM

To validate *in silico* model prediction with phenotypic observations, the growth (respiration) phenotype of three representative APEC isolates, chosen from phylogroups B2, C and G, were tested using the Biolog Phenotypic Microarray analysis (PM1, PM2A: carbon source plates, and PM4A: sulphate and phosphate plate). The respiration measurements are available in Supplementary Material 2, Table S3. In total, 125 nutrient substrates were analysed in the APEC model, and 119 substrates were analysed in *i*jO1366 (Supplementary Material 2, Table S4). The Pearson correlation coefficient of all three comparison experiments showed mostly positive correlations; the *i*JO1366a model showed a weak positive correlation (r^2^ values between 0.07 and 0.17, Supplementary Material 1, Figure S3), while the *i*JO1366b and APEC models showed a moderate positive correlations (r^2^ values between 0.50 and 0.59, Figure 6). Under the optimised media settings, the APEC model showed an improved performance with the representative phylogroups C (r^2^=0.59, *p* < 0.005) and G (r^2^=0.59, *p* < 0.005), while it showed decreased performance with phylogroup B2 (r^2^=0.50, *p* < 0.005), compared to model *i*JO1366 (Figure 6).

**Figure 6.**
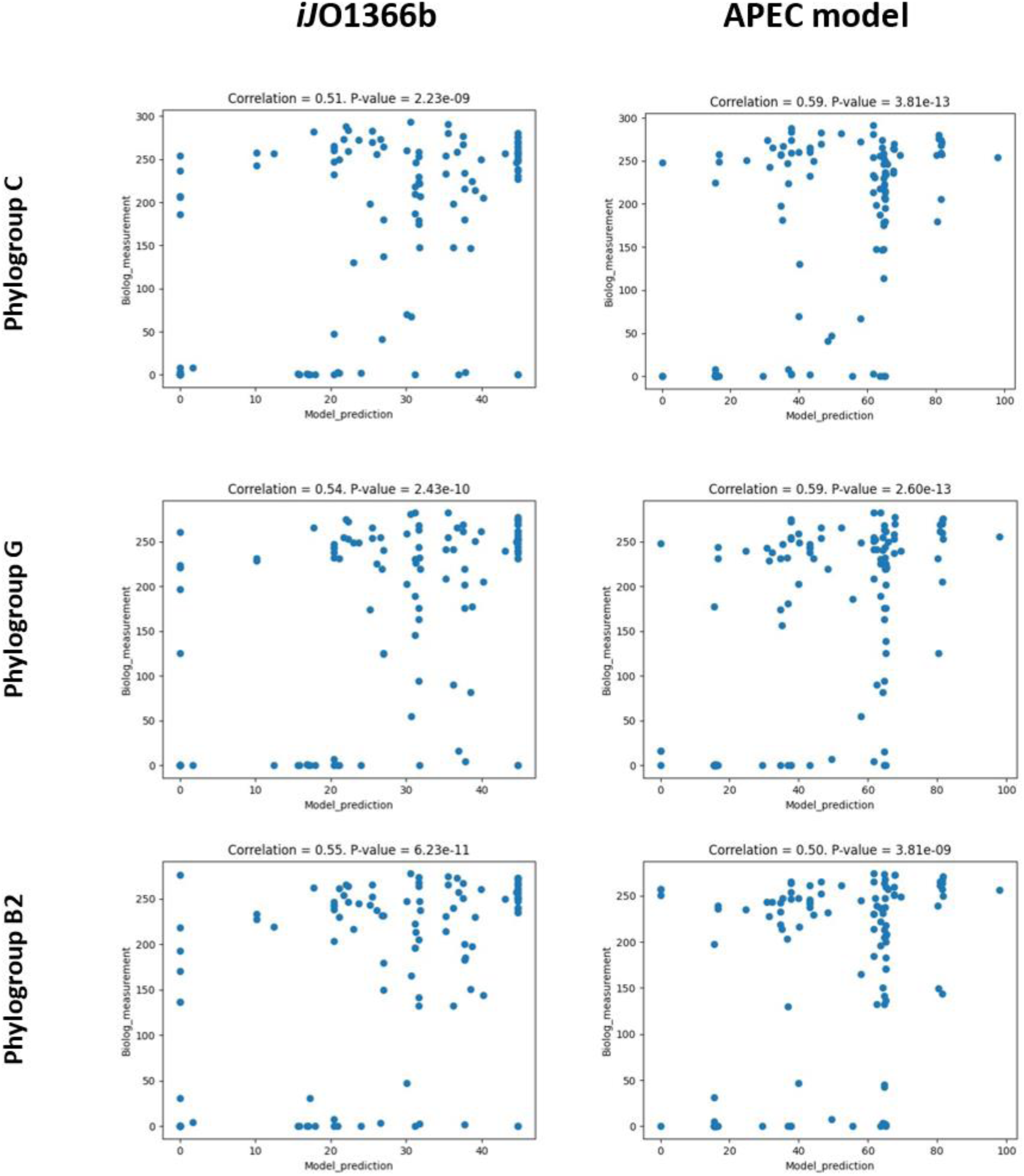
Scatter plots comparing computational predictions from the *E. coli* K-12 *i*JO1366 model and the APEC model for three APEC isolates representing phylogroups B2, C and G, with Biolog phenotypic observations with single nutrient source utilisation. The APEC isolates used are from phylogroup C (SAP0009), G (SAP0545) and B2 (SAP0474). The scatter plots show the model prediction (X-axis) using *i*JO1366b (Left) and APEC model (Right) and phenotypic observation using Biolog measurement (Y-axis). The correlation value shown above graphs is the Pearson correlation coefficient r^2^ value.

### Comparison of the APEC GEM with phylogroup-specific sub-models

We also explored the performance of phylogroup-specific APEC model based sub models. The default media setting of phylogroup based sub-models were modified from the APEC GEM, with the import of Fe(II)-Nicotianamine (Fe(II)-NA or Fe^2+^-NA) added as it was required for model performance of sub-models for phylogroups C and G. In total, *in silico* flux balance analysis (FBA) of 142 nutrient substrates was performed with phylogroup-specific sub-models and then compared to growth simulation of APEC GEM according to Biolog data (Supplementary Material 2, Table S4). In Figure 7, the relationship between the computational prediction from the sub-models and their Biolog measurements showed moderate positive correlation (r^2^ values of 0.51, 0.64 and 0.3 for phylogroups C, G and B2, respectively). Compared to the computational prediction by the pan model APEC GEM, the sub-models of phylogroup C performed less well under similar media input, while the phylogroup G and B2 sub-model showed improved performance compared to APEC model.

**Figure 7.**
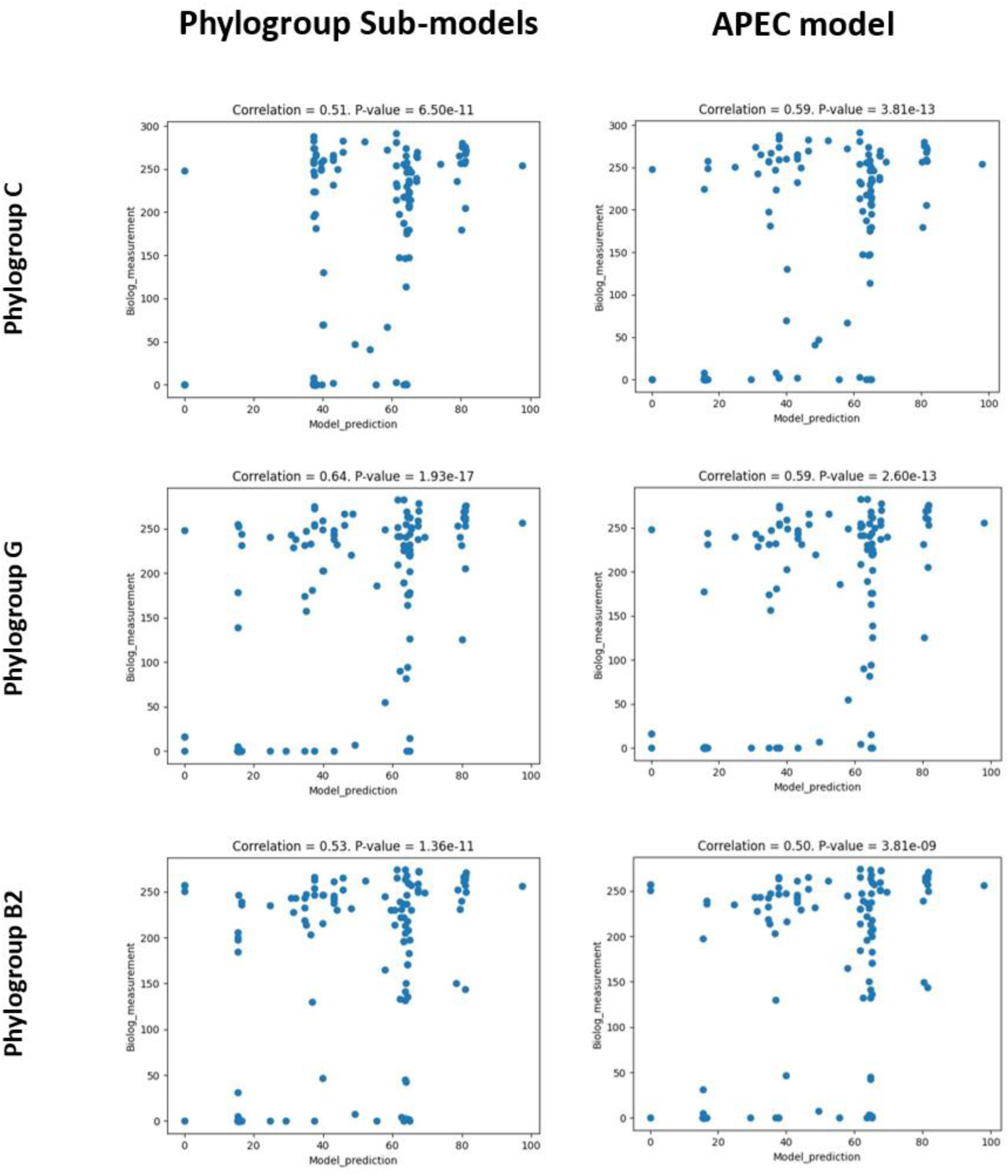
Scatter plots comparing computational predictions from the APEC phylogroup-specific sub-models for phylogroups B2, C and G with the general APEC model for three APEC isolates representing phylogroups B2, C and G, with Biolog phenotypic observations with single nutrient source utilisation. The APEC isolates used are from phylogroup C (SAP0009), G (SAP0545) and B2 (SAP0474). The scatter plots show the model prediction (X-axis) using *i*JO1366b (Left) and APEC model (Right) and phenotypic observation using Biolog measurement (Y-axis). The correlation value shown above graphs is the Pearson correlation coefficient r^2^ value.

### Identification of 3-hydroxyphenylacetate as a phylogroup C APEC-specific nutrient

During querying of the phylogroup-specific APEC GEMs, one of the reactions predicted to be specific for phylogroup C APEC isolates was associated with the metabolism of 3- and 4-hydroxyphenylacetic acid:

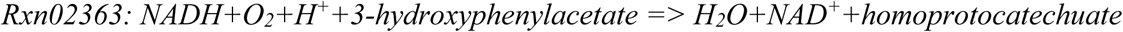

This reaction is catalysed by 4-hydroxyphenylacetate 3-monooxygenase oxygenase, encoded by gene *hpaB* (4-hydroxyphenylacetate 3-monooxygenase, EC 1.14.13.3). The *hpaB* gene was present in all phylogroup C (n=21/21) APEC, while absent in phylogroup B2 (n=0/31) and G (n=0/19) APEC. The *hpaB* gene was detected in phylogroups A and B1 as well, but these constitute minor phylogroups for APEC. The metabolic variation on 3-HPAA metabolism was measured using M9 minimal medium. As is shown in Supplementary Material 1, Figure S4, five phylogroup C isolates were all able to grow with the three different concentrations of 3-HPAA tested, whereas six phylogroup B2 and two phylogroup G isolates were unable to grow with 3-HPAA as the single carbon source (Supplementary Material 1, Figure S5), but all grew in LB medium.

The metabolism of 3-HPAA in *E. coli* has been described previously [43, 44]. However, the intracellular/extracellular reaction of 3-HPAA in *E. coli* has not yet been documented in the ModelSEED database. Therefore, we manually added the exchange reaction of 3-HPAA into the sub-models, in order to perform the growth simulation with/without supplementary of 3-HPAA. The additional reactions are presented as below (cpd03320 is the compound ID of 3-HPAA in APEC GEM and relative sub-models, *e* represent for extracellular; *b* represents for periplasm; *c* represent for cytoplasm):

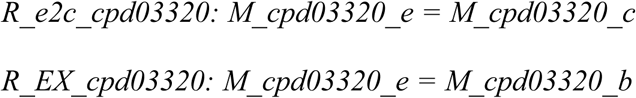

The lower bound (lb) of the 3-HPAA exchange reaction was adjusted to 0, -10, -100 and -1000 in the APEC GEM sub-model B2, C and G, to simulate APEC growth in response to four different concentrations of 3-HPAA. The growth prediction of different sub-groups and M9 experimental observation are summarised in Figure 8. In sub-models B2 and G, the relatively minimal biomass (∼15) was produced with the lower bound of 3-HPAA exchange reaction adjusted to 0, -10, -100, and -1000 (increasing uptake), which was assumed as no growth for the modelling purpose. This agreed with the M9 minimal media experimental data, where isolates from phylogroup B2 and G were unable to use 3-HPAA as the sole carbon source. However, growth was predicted in sub-model C, matching the setting of increased uptake of 3-HPAA (Figure 8).

**Figure 8.**
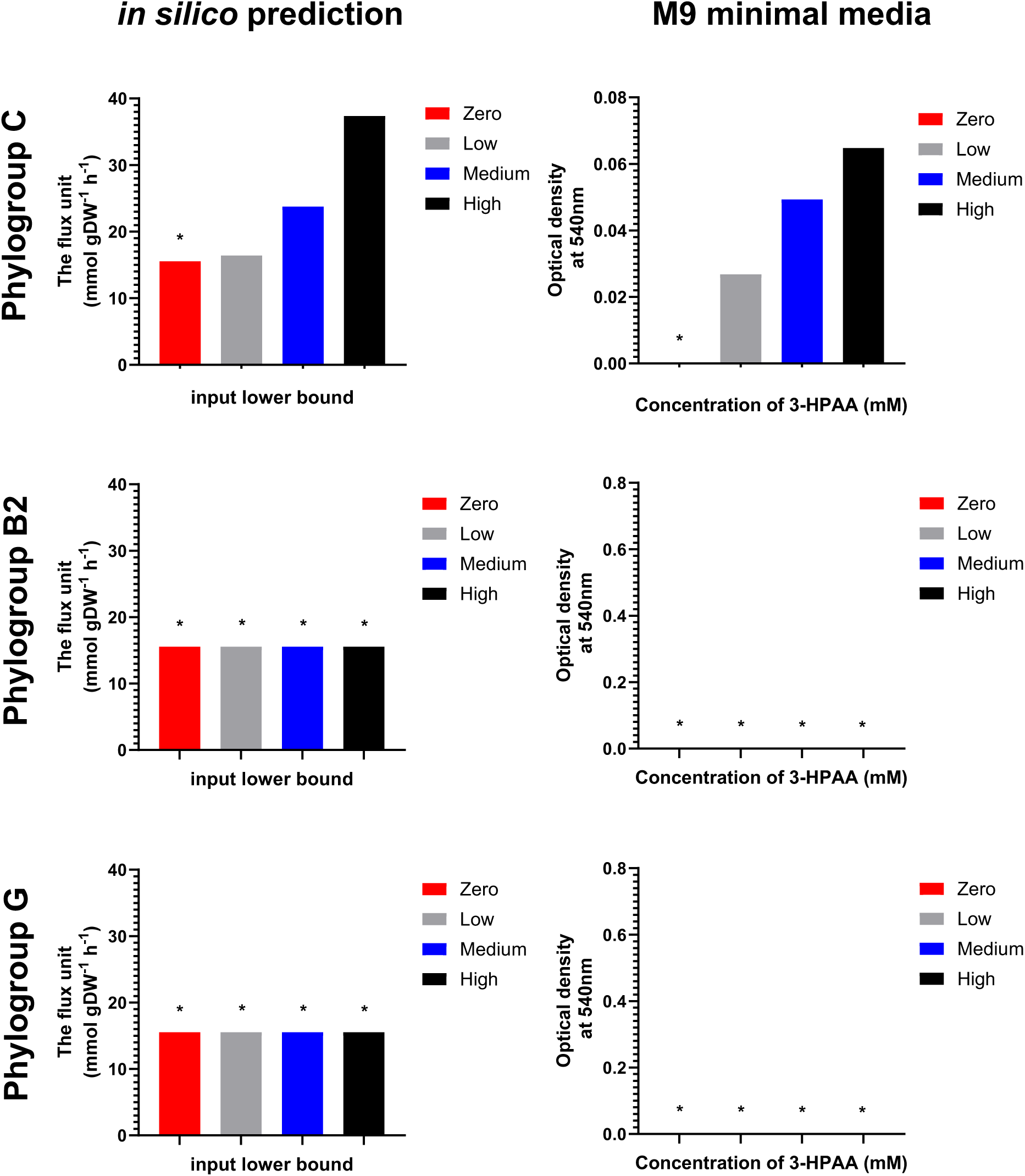
The comparison between *in silico* prediction and M9 minimal medium measurements of different phylogroup APEC isolates on utilisation of different concentration of 3-hydroxyphenoylacetic acid (3-HPAA). The left figures show the *in silico* prediction using phylogroup-based sub-model (C, G and B2) under the constrain of 3-HPAA exchange reaction lower bound adjusted to 0, -10 (low), -100 (medium) and -1000 (high), respectively (-1000 refers to unlimited uptake of inputs). The right figures show the phenotypic measurements in M9 minimal media with supplement of 0 mM (zero), 0.5 mM (low), 0.75 mM (medium), and 1 mM (high) 3-hydroxyphenoylacetic acid as sole carbon source. The measurements indicate the mean of three biological endpoint (t=18h) readings. More detailed growth curves are provided in Supplementary Material 1, Figures S4 and S5. Asterisks indicate an absence of growth.

## DISCUSSION

In this study, we have investigated the metabolic capabilities of a representative set of 114 avian pathogenic *E. coli* isolates representing different phylogroups. A genome scale metabolic model was constructed based on the genome sequences of these APEC isolates and used to 1) perform growth simulation experiments which were validated using phenotypic assays 2) perform gene essentiality analysis 3) investigate functional difference between phylogroups and develop phylogroup based sub-models.

The comparative genomic analysis performed in this study confirmed that the APEC pathotype is genetically variable, with phylogroups B2, C and G being the dominant lineages [4, 5, 45]. The resulting genome scale metabolic profile of the 114 APEC isolates (Figure 3, Supplementary Material 2, Table S9) showed strong conservation in central metabolism involved in energy production, with the ratio of core metabolism versus accessory metabolism higher than in previous studies, where multiple *E. coli* pathotypes and commensal isolates were involved [9, 46]. The major metabolic variances between isolates are associated with metabolism of carbohydrates and amino acids, as well as exchange/transport reactions. This was confirmed using Biolog metabolic analysis performed in this study. As shown in Figure 3 and Supplementary Material 2, Table S9, the phylogroup B2 isolates showed distinct metabolic features (mostly missing metabolic reactions) compared to others, which was previously also observed [36, 47, 48]. This is consistent with the view that loss and acquisition of genes during evolution differentiates isolates into different lineage with unique metabolic profiles and adaptation to specific niches.

When comparing the predictions of the APEC model with the Biolog phenotypic measurements, the association between APEC GEM prediction and phenotypic observations was moderately positive (r^2^ values between 0.5 and 0.6). The vast numbers of reactions and genes encompassed within metabolic models potentially leads to a high risk of gaps or errors within the metabolic network. Furthermore, the reconstruction of GEMs is severely limited by the paucity of verified metabolic pathways that are specific to APEC and current knowledge in our database, including errors introduced by automatic annotation of genome sequences in the respective databases. From the perspective of APEC diversity, the pan-genome scale metabolic model is too extensive to represent the differences and specifics of certain APEC lineages. Notably, in this study, we used the Biolog platform as the indicator of biomass formation of APEC isolates in response to different nutrient sources, while this assay is based on the measurement of microbial respiration and reflected by dye changes. These two different measurements may lead to false-positive or -negative results during the coefficient relationship analysis. This was addressed by changing the function from biomass production to indicator dye reduction used in the Biolog plates, which leads to a significantly decrease in false negative results [8]. Alternatively, a comprehensive screening of isolates using different nutrient sources by M9 minimal medium may provide more direct observation for model validation. Typically, the phenotype of the Keio collection was used in model validation of *i*JO1366 [14]. However, this was not appropriate for our study due to the significant differences between the laboratory strain K-12 and the APEC pathotype.

Moreover, the APEC pathotype has a diverse metabolic profile and construction of a single gene knockout library representing the APEC pathotype is not possible, which made it difficult to apply the APEC metabolic model for the pathotype. However, the APEC metabolic model provides a comprehensive database of metabolic capabilities for a diverse panel of APEC, and as such can be used for initial predictions, and can be used as the basis for the construction of lineage-based strain sub-models, as performed here for the dominant phylogroups B2, C and G.

Although no significant improvement in the association strength between computational prediction of sub-models and the phenotypic measurement of the corresponding APEC isolates were observed, the APEC model showed its potential and plasticity as a pan model used for the construction of phylogroup B2, C and G sub-models. The phenotypic difference in catabolism of 3-HPAA in phylogroup B2, C and G were successfully predicted by those phylogroup based sub-models as a proof of principle. Interestingly, 3-hydroxyphenylacetate and 4-hydroxyphenylacetate are the main phenolic acids derived from flavonoid quercetin (which is commonly added to poultry feed) in the colorectum [49, 50]. Therefore, these compounds are likely to accumulate in the avian intestine and its utilisation could be a hypothetical nutrient advantage in the intestinal microbial community. The phylogroup C sub-models not only predicted the growth of phylogroup C APEC isolates using 3-hydroxyphenylacetate as sole carbon source, but also reflected the biomass increases alongside the enhancing lower bound setting of 3-hydroxyphenylacetate (Figure 8, Supplementary Material 1, Figure S4).

The models generated in this study can be further exploited for the identification of additional pathways that are unique to APEC lineages. Although APEC are extraintestinal pathogens, potential APEC isolates are present in the intestinal *E. coli* population [7], and may form a reservoir for future infections within flocks. These models may inform the targeted suppression of such APEC populations thereby benefitting avian health and welfare in the long term. The approach taken may also be applicable for other genetically diverse intestinal or extraintestinal pathogens dispersed over multiple lineages. Furthermore, in the future, the rapid progress in software development (e.g. Bactabolize [51]), availability of APEC genome sequences and associated repositories may expedite the generation of genome scale metabolic models to revisit or improve existing metabolic models for APEC and other *E. coli* pathotypes.

## ACKNOWLEDGMENTS

The authors would like to acknowledge the diagnostic laboratories for providing the isolates used in the study.

## Abbreviations

SRA: Sequence Read Archive
APEC: Avian Pathogenic *Escherichia coli*
ExPEC: Extra-Intestinal Pathogenic *E. coli*
GEM: Genome scale metabolic reconstruction
MR: antimicrobial resistance
AMP: ampicillin; Cm, chloramphenicol
CAT: chloramphenicol resistance cassette
PCR: Polymerase Chain Reaction
RT-qPCR: reverse transcription quantitative-PCR
3-HPAA: 3-hydroxyphenylacetate
SBC: stoichiometrically balanced cycles
BCR: biomass consumption reactions
FBA: flux balance analysis.

## Notes

**Conflict of interest.** The authors declare that there are no conflicts of interest.

### Competing Interest Statement

The authors have declared no competing interest.

### Summary of Updates

Revision based on comments from peer reviewers. Some additional text in the manuscript, more information added to Table S1

