## Supplementary Figures S1-S5 for "Use of Genome Scale Metabolic Reconstructions of Avian Pathogenic *Escherichia coli* (APEC) phylogroups for the identification of lineage-specific metabolic pathways"

Supplementary Figures 1-5 for:

**Use of Genome Scale Metabolic Reconstructions of Avian Pathogenic *Escherichia coli* (APEC) phylogroups for the identification of lineage-specific metabolic pathways**

Huijun Long, Jai W. Mehat, HuiHai Wu, Arnoud H. M. van Vliet, Roberto M. La Ragione

**Figure S1.** Summary of the deleted reactions for construction of phylogenetic group specific sub-models based on APEC model.

**Figure S2.** The distribution of essential genes predicted by *in silico* analysis against glucose and glycerol as sole carbon sources, using the APEC model in different functional categories.

**Figure S3.** Scatter plots comparing computational predictions from the *E. coli* K-12 *i*JO1366 model and the APEC model for three APEC isolates representing phylogroups B2, C and G, with Biolog phenotypic observations with single nutrient source utilisation.

**Figure S4.** Growth of five representative Phylogroup C APEC isolates in M9 minimal media supplemented with 3-hydroxyphenylacetic acid (panels A-E) and in LB broth (panel F), respectively.

**Figure S5.** The growth curve of phylogroup B2 and G APEC isolates in M9 minimal media supplemented with 3-hydroxyphenylacetic acid (1 mM, 0.75 mM and 0.5 mM) and in LB broth, respectively.

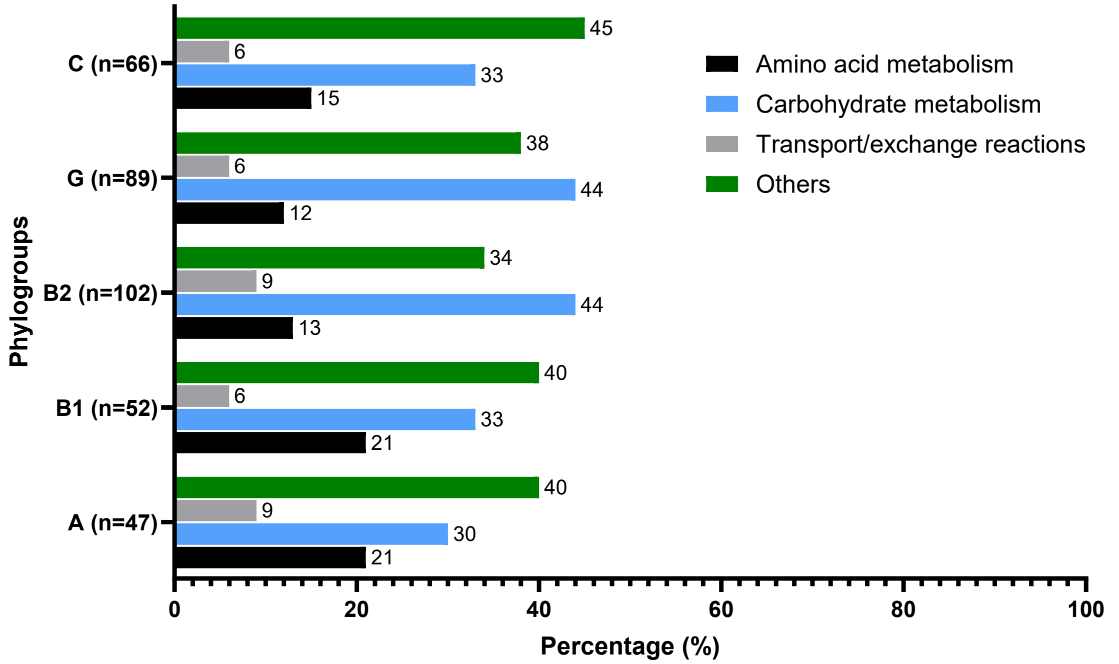


**Figure S1.** Summary of the deleted reactions for construction of phylogenetic group specific sub-models based on APEC model. A total of 133 reactions were removed in construction of phylogroups-specific sub-models for 114 APEC isolates in this study. Colour code of functional categories of reactions: Amino acid metabolism = Black; Carbohydrate metabolism = Blue; Transport/exchange reactions = Grey; Others* = Green. *Other reactions including Cell wall/ membrane/ envelope metabolism, Cofactor and prosthetic group metabolism, Energy production and conversion, Inorganic ion transport and metabolism, Lipid metabolism, Nucleotide metabolism and other described in Figure 2.

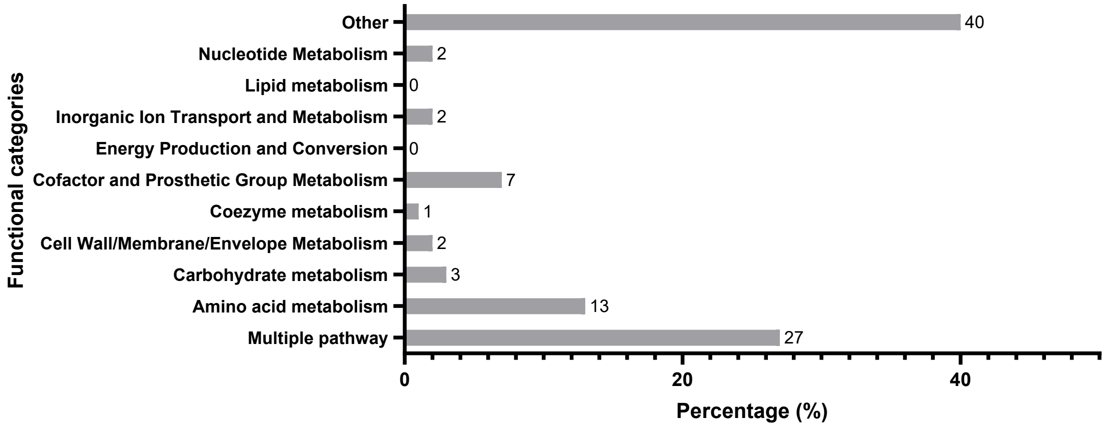


**Figure S2.** The distribution of essential genes predicted by *in silico* analysis against glucose and glycerol as sole carbon sources, using the APEC model in different functional categories.

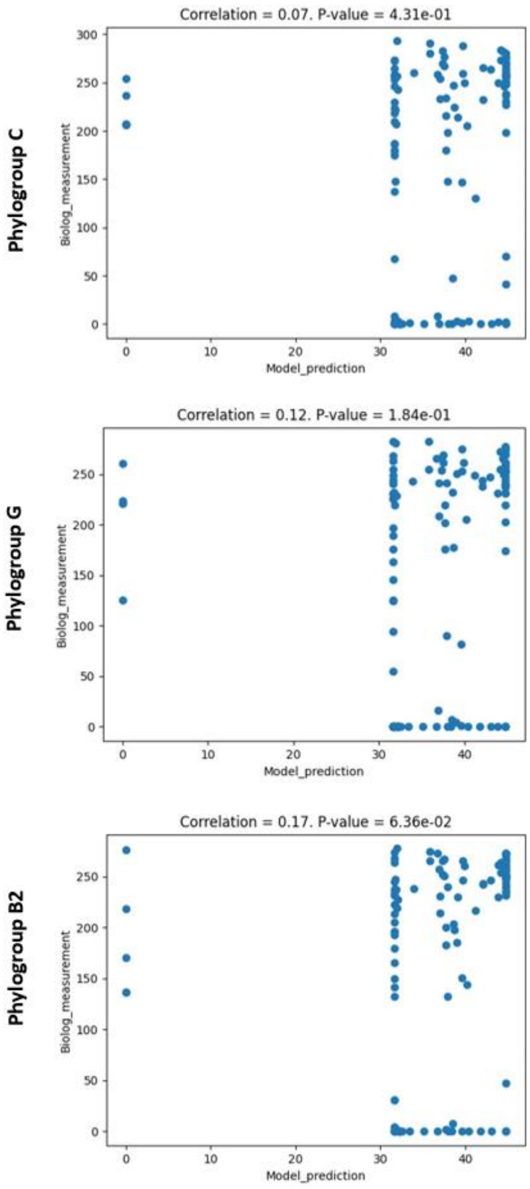


**Figure S3.** Scatter plots comparing computational predictions from the *E. coli* K-12 *i*JO1366 model and the APEC model for three APEC isolates representing phylogroups B2, C and G, with Biolog phenotypic observations with single nutrient source utilisation. The APEC isolates used are from phylogroup C (SAP0009), G (SAP0545) and B2 (SAP0474). The scatter plots show the model prediction (X-axis) using *i*JO1366a (Left) and APEC model (Right) and phenotypic observation using Biolog measurement (Y-axis). The correlation value shown above graphs is the $r^2$ value.

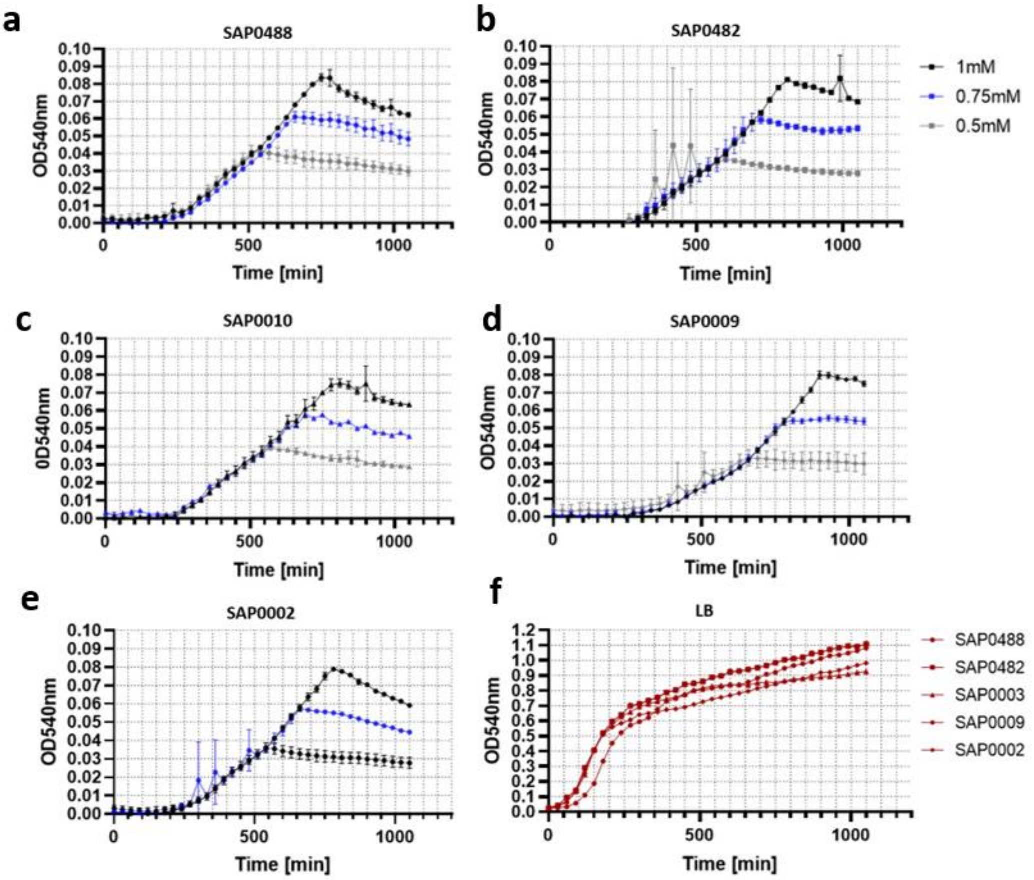


**Figure S4.** Growth of five representative Phylogroup C APEC isolates in M9 minimal media supplemented with 3-hydroxyphenylacetic acid (panels A-E) and in LB broth (panel F), respectively. The growth curves show the mean of 3 biological replicates, using optical density measurements at 600 nm wavelength for every 30 min in 18h, with error bars indicating the standard error of the mean (SEM). Panels A-E illustrates the growth curves of phylogroup C APEC isolates in M9 minimal media supplemented with three concentrations of 3-hydroxyphenylacetic acid (1 mM, 0.75 mM and 0.5 mM). Panel F shows growth curves of all group C isolates tested in this study in LB broth.

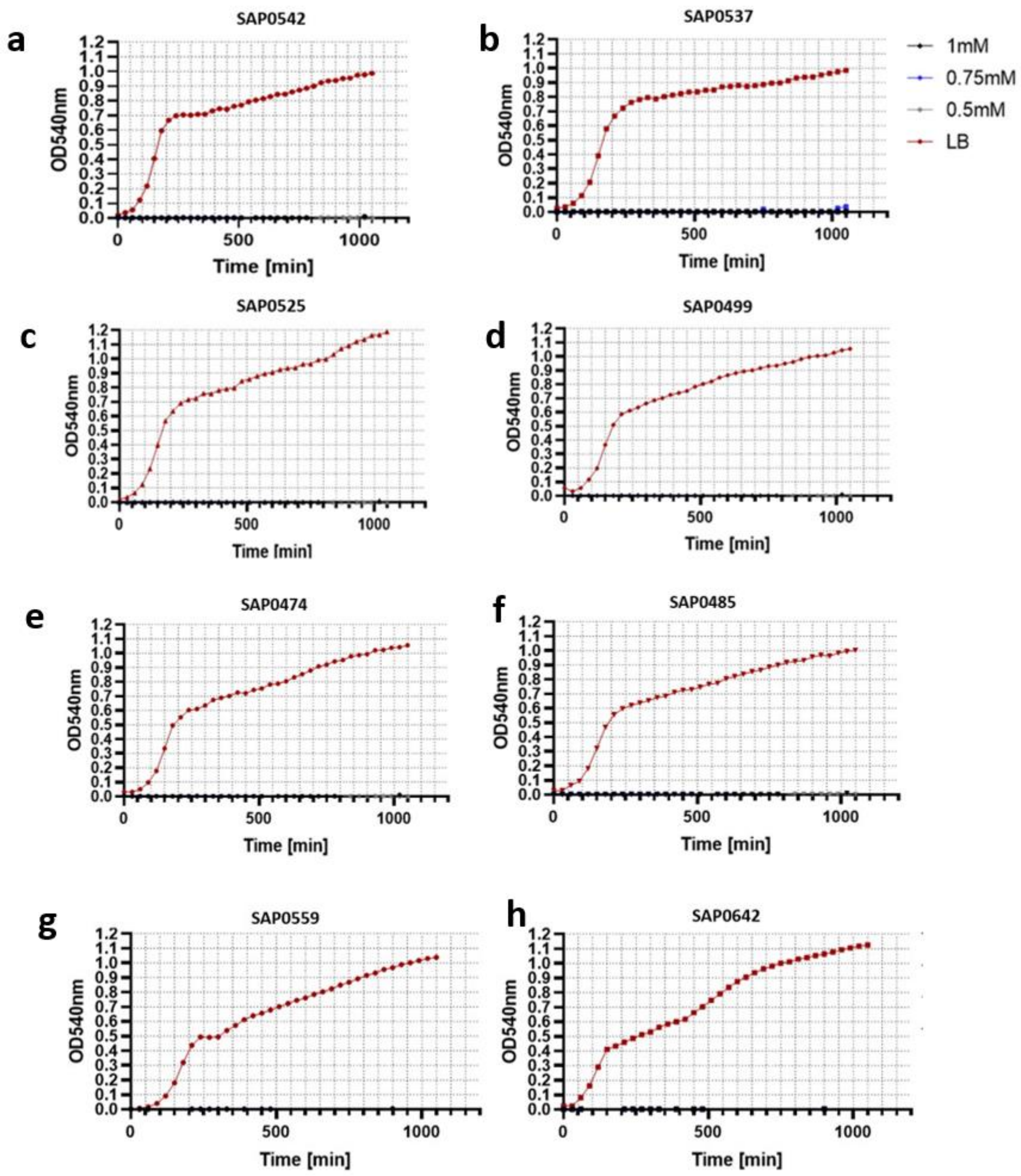


**Figure S5.** The growth curve of phylogroup B2 and G APEC isolates in M9 minimal media supplemented with 3-hydroxyphenylacetic acid (1 mM, 0.75 mM and 0.5 mM) and in LB broth, respectively. The growth curves show the mean of 3 biological replicates, using optical density measurements at 600 nm wavelength for every 30 min in 18h, with error bars indicating the standard error of the mean (SEM). Panels A-F shows the phylogroup B2 APEC isolates used in this study whilst panels G-H shows the phylogroup G APEC isolates used in this study. There was no growth observed with any of the phylogroup B2 and G isolates with any of the concentrations of 3-HPAA added.
